# Brain–heart dynamics during emotional processing under uncertain conditions: An index of depression risk

**DOI:** 10.1101/2021.12.06.471520

**Authors:** Hui-Ling Chan, Noriaki Kanayama, Ryohei Mizuochi, Shigeto Yamawaki, Maro G. Machizawa

**Author notes:** Corresponding authors: Hui-Ling Chan and Maro G. Machizawa. 62561.

## Abstract

Recent studies have highlighted the essential role of interoception in healthy emotional processing and the pathology of major depressive disorder. However, it is unclear how individual differences in healthy people with high depression risk (HDR; i.e., individual differences in depression risk) are related to the neurophysiological underpinnings of interoception and emotional reactions under different degrees of certainty. We examined whether an individual’s depression risk mediates the relationships of heartbeat-evoked potentials (HEPs), a neurophysiological marker of cardiac interoception, with heart rate (HR) and heart rate variability (HRV), indices for cardiac reactivity. In a concurrent electroencephalogram-electrocardiogram (EEG-ECG) experiment, 26 healthy participants completed an emotion-evoking picture-evaluation task. Each trial began with a differential auditory cue that was associated with the certainty of subsequently seeing a pleasant or unpleasant picture. The results showed the following: after participants saw a cue of uncertainty, HDR was associated with 1) reduced neural activity when anticipating upcoming pictures and 2) enhanced physiological reactions to <u>unexpected</u>, unpleasant pictures. These results suggest that weaker predictions and augmented prediction errors for negative emotional stimuli reflect depression risk. Moreover, depression risk significantly mediated the relationships between HEPs and HR and between HEPs and HRV for unexpected, unpleasant stimuli. This study provides evidence that interoception and autonomic cardiac regulation may be altered by depression risk. This highlights the insights provided by specific indices of brain–heart interactions, such as HEPs, into the underlying activity of the autonomic nervous system and unique interoceptive disturbances associated with depression risk.

## 1. Introduction

Depression, an emotional state marked by a consistently low mood, is characterized by altered emotional processing (1). Efforts to understand emotional processing have yielded evidence that suggests that interoception plays a crucial role (2). Impairments in interoceptive function in depression have been widely explored (3–5); these include decreased heartbeat counting accuracy (4), altered activity of brain regions supporting interoception (6, 7) and disturbed visceral activity (8, 9). All these findings suggest that further investigations of the interoceptive processing underlying emotional processing in depression are needed.

Moreover, individuals with high depression risk (HDR), i.e., an elevated risk of developing major depressive disorder (MDD) (10), also show altered interoception, such as reduced connectivity in the brain’s interoceptive network (11, 12). Thus, alterations in the interoceptive system may occur along a continuum, as do levels of trait depression. However, the link between the interoceptive mechanisms underlying emotional processing and depression risk remains unknown.

Interoception, as narrowly defined, is a bottom-up process that involves the perception of bodily states. These bodily signals influence brain functions and thus shape sensory perception, emotion, and cognition (13–15). Accumulated evidence suggests that interoception includes the integration of both bottom-up (stimulus-based) and top-down (model-based) interoceptive information (16, 17). The integration of the current sensory experience and the prediction from the internal model allow the calculation of interoceptive prediction errors (18), which are followed by bodily responses or updates of the internal model. The disturbance of interoception in MDD may result from an inefficient internal model and unreliable prediction errors (19). Using the structural equation model, a study demonstrated that low levels of interoceptive awareness lead to alexithymia and emotional dysregulation, which in turns predict depression risk (20); this study used a self-report questionnaire to assess interoceptive awareness, namely, the Multidimensional Assessment of Interoceptive Awareness-2 (MAIA-2) (21). In contrast to the aforementioned study, which collected subjective questionnaire data using the MAIA-2, we recorded objective neurophysiological data. To further investigate the link between interoceptive processing and depression risk, we assessed the disturbance of interoceptive responses along with depression risk (via a subjective report) based on neural and physiological responses to predictable and unpredictable emotional events.

The cardiac autonomic nervous system (ANS) automatically responds to emotional events. For example, heart rate (HR) (22) and heart rate variability (HRV) (23) are well-known markers for cardiac reactivity that reflect changes in valence (pleasant or unpleasant emotional states). Emotional events also influence heartbeat-evoked potentials (HEPs) (24), which are the *neural* activity corresponding to each heartbeat, as well as the correlation between HEPs and HRV (25). HEPs reflect interoceptive cardiac processing in the brain (26). The generation of HEPs has been suggested to reflect the projection of cardiac *afferent* signals from the heart to brain regions, such as the insula (27). Therefore, HEPs and HR/HRV are putatively excellent candidates for examining the neural and cardiac substrates underlying emotional processing, respectively.

The interoceptive system is responsible for regulatory processes that maintain or restore homeostatic balance at both the conscious and unconscious (autonomic) levels (28, 29). A study examining HRV during neurofeedback training revealed an inverse correlation between HRV and HEPs to during emotional conditions (25). Moreover, in a magnetoencephalography study (30), subjects’ heartbeat-evoked responses not only were affected by hearing their own name but also led to bias when making judgments of the subject’s own name. Their results suggested that heartbeat-evoked responses have a reciprocal causal relationship with prestimulus predictions and poststimulus evaluations. Overall, the cardiac interoceptive system may interact with both cardiac signals and external stimuli. Alterations in the relationship between HEPs and cardiac reactivity in response to external stimuli may reflect dysfunction of brain–heart interaction. Compared to healthy participants, MDD patients have reduced HEPs during interoceptive tasks (31) and reduced resting-state HRV (32). Individuals with HDR also show lower HR recovery rates in response to unpleasant images (33) and lower resting-state HRV (34) than individuals with low depression risk. While the literature suggests that depression is linked to dysfunction of the interoceptive system and altered cardiac reactivity, the effects of depression risk on brain–heart interactions and the link between the cardiac interoceptive system and cardiac reactivity remain unclear.

The present study investigated the neural substrates of the cardiac interoceptive system and cardiac reactivity to emotional events under conditions of certainty and uncertainty in healthy individuals. Both electroencephalography (EEG) and electrocardiography (ECG) were used to monitor the participants while performing an emotion-evoking picture-evaluation task. A precue tone was played to induce anticipatory responses, followed by the display of either a pleasant or an unpleasant image. HEPs were assessed as an index of neurophysiological interaction, and HR and HRV were assessed as indices of cardiac reactivity.

Previous studies have shown that compared to conditions of certainty, conditions of *uncertainty* induce greater regulatory responses in both the cardiac (35, 36) and neuro-cardiac (37, 38) domains, which may result from large prediction errors. Abnormal anticipation of future events in the central interoceptive system has been observed in individuals with HDR (39, 40). Based on previous studies, we hypothesized that stimuli with *negative* valence (i.e., unpleasant pictures) would elicit larger responses under conditions of *uncertainty*. A mediation analysis was performed to further examine the role of depression risk in cardiac regulation; specifically the interactions between the neural substrate of the cardiac interoceptive system and physiological reactivity were analyzed. Since little is known about the spatiotemporal dynamics of HEPs, we further examined which specific temporal and spatial features of HEPs play pivotal roles in the relationships among the three factors (cardiac reactivity, neuro-cardiac reactivity, and depression risk), in a data-driven manner. Overall, we examined how cardiac and neuro-cardiac signals interact with certainty and the emotional processing associated with depression risk.

## 2. Materials and Methods

### 2.1. Participants

Twenty-six healthy participants (16 females; mean ± standard deviation (*SD*) of age: 22.19 ± 1.86 years) with no history of psychiatric, psychological, or cardiac disorders participated in an emotion-evoking picture-evaluation task with concurrent EEG and ECG recording. The data were acquired as part of a previous study (41). Participants were recruited locally, and written informed consent was obtained from participants in person prior to the experiment, in line with approval of the local ethical committee (Epidemiological Research Ethics Review Committee of Hiroshima University; approval number: E-Epidemic-0965-1; August 12^th^, 2014). The recruitment period for the experiment was from 26 November 2014 to 7 April 2015.

### 2.2. Questionnaire procedure

The level of depression risk was evaluated using the Japanese version of the Beck Depression Inventory-II (BDI) (42, 43). The overall mean and *SD* of the BDI score were 5.65 and 7.22, respectively. The distribution of BDI scores was left-skewed, as shown in Fig 1. Kolmogorov‒Smirnov two-sample tests performed on the distributions of this dataset and a public dataset of 121 healthy participants collected by Cavanagh (44) indicated that the distribution of BDI scores in our dataset was not significantly different from that of the large-sample dataset (*p* = .19, n.s.). Extreme group approaches are suitable for analyzing and visualizing skewed data or data with small sample sizes (45) and avoiding misclassification of samples with scores near the median (46). For display purposes, in this study, participants were split into two groups using a quartile split (47). Participants with BDI scores higher than the third quartile cutoff (BDI scores ≥ 5) or lower than the first quartile cutoff (BDI scores ≤ 2) were categorized into high BDI (n = 8, BDI scores = 14.12 ± 7.42) or low BDI (n = 13, BDI scores = 1.23 ± 0.89) groups, respectively. Notably, these groups were used for the sake of visualization only and were not included in statistical analyses.

**Fig 1.**
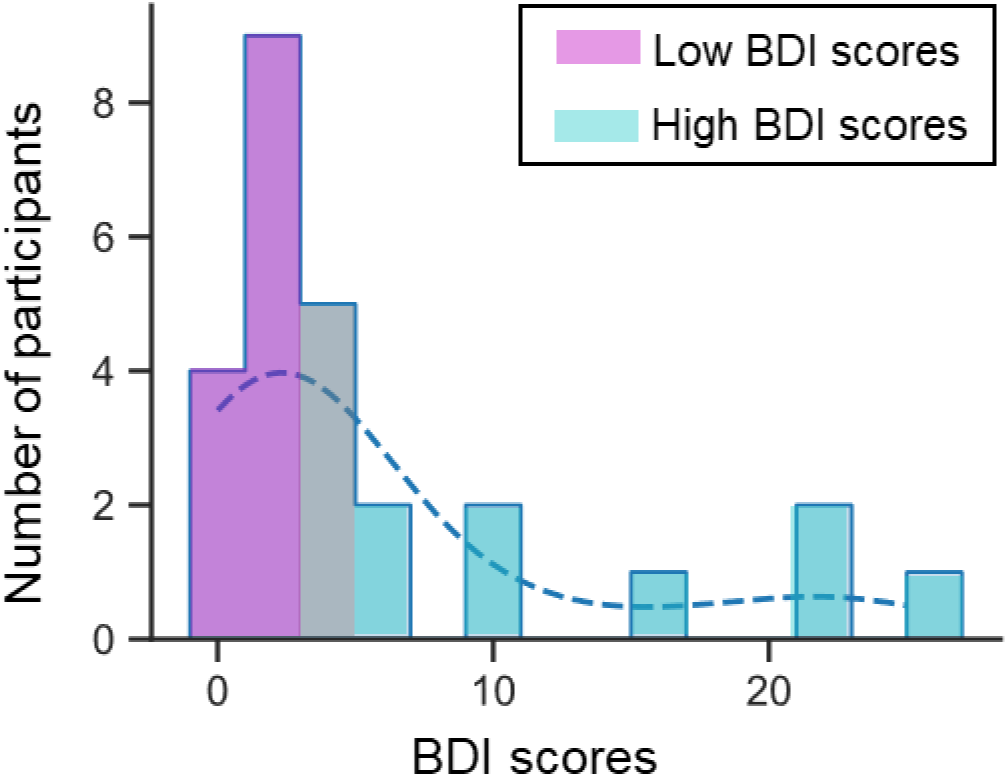
Histogram showing the distribution of the BDI scores of all participants. The dashed line represents the kernel density estimate, showing the smoothed distribution of BDI scores. Cyan and magenta colored bars indicate the high BDI group and low BDI group, respectively. The gray color indicates participants with BDI scores between the thresholds for assignment to the two groups. This quartile split was used to visualize the results of further analyses. BDI: Beck Depression Inventory.

### 2.3. Behavioral task and procedure

The emotion-evoking picture-evaluation experiment consisted of 240 trials. Each trial comprised a precue and picture-presentation phase, followed by a subjective evaluation of valence. During the precue (‘cue’ phase), an auditory tone was presented for 0.25 s, followed by a 3.75-s delay. During the picture-presentation phase (‘pic’ phase), a pleasant or an unpleasant picture was displayed for 4 s, during which time the participants were asked to evaluate the emotion they felt. Participants rated the valence of their emotional responses on a four-point Likert scale (from most unpleasant to most pleasant) during the evaluation phase, immediately after seeing the picture. Three different tones (low, mid, and high at 500, 1,000, and 1,500 Hz, respectively) were used as auditory cues to indicate one of three certainty conditions: predictably pleasant (PP), unpredictable (UX), and predictably unpleasant (PU). The unpredictable condition was further separated into two subconditions based on the picture’s valence: unpredictably pleasant (UP) and unpredictably unpleasant (UU). Note that the letter X is used in this manuscript to indicate two possible conditions. For example, UX represents both the UP and UU conditions (unpredictable conditions, regardless of the valence of the subsequent picture), whereas XU represents both the PU and UU conditions (negative picture conditions, regardless of the tone). The mid tone (1,000 Hz) was the cue for the UX condition. The 500- and 1,500-Hz tones were used for the PP or PU conditions (the assignment was counterbalanced across participants). Each of the three certainty conditions consisted of 80 trials. In the UX condition, 40 trials involved pleasant pictures (UP), and the other 40 involved unpleasant pictures (UU). Pictures were selected from the International Emotional Picture System (IAPS) (48). Finally, the participants rated their emotions after seeing the images. Further details have been published elsewhere (41).

### 2.4. EEG-ECG recordings

During the task, brain and cardiac activity were simultaneously recorded using the BioSemi ActiveTwo system (BioSemi, Amsterdam) at a sampling rate of 2,048 Hz. The 64-channel EEG electrodes were placed according to the conventional international 10-20 system. In addition to EEG channels, ECG electrodes were placed on the left wrist and right side of the back of the neck to monitor bipolar ECG signals. Additionally, conventional bipolar electrooculography (EOG) was used to measure vertical and horizontal eye movements. Four EOG electrodes were individually placed above and below the right eye and to the left and right of the outer canthi. An additional channel was placed at the tip of the nose for offline rereferencing.

### 2.5. HR and HRV computation

ECG signal preprocessing, R peak detection, and the calculation of instantaneous heart rate (HR) were conducted using the Python packages BioSPPy (49) and NeuroKit (50). Raw ECG signals were first bandpass-filtered with a 3–45 Hz finite impulse response (FIR) filter, followed by detection (51) and location correction of R peaks by relocating each R peak to the time point with the maximum ECG signal within the time range of -50 to 50 ms. Instantaneous HR was calculated as the inverse of the interbeat interval (IBI) between two successive R peaks. For subsequent analyses, instantaneous HR data points exceeding the physiological range (40–200 bpm) were discarded. Finally, HR time series were obtained by applying boxcar smoothing with a window size of 3 and cubic spline interpolation to the instantaneous HR values.

We used the *SD* of the normal-to-normal IBI (SDNN) as an HRV index in the current study (52–54). To obtain the HRV time series, IBIs with z scores larger than 1.96 were discarded, and then the remaining IBIs were upsampled to 4 Hz using cubic spline interpolation. Since accurate measures of SDNN can be obtained from 10-s recordings (55, 56), we then applied a 10-s sliding window with a 0.25-s step size to the interpolated IBIs to compute the *SD* of IBIs within the window as the instantaneous HRV value at the center of the window. The resulting HRV time series (with a 4-Hz sampling rate) was further upsampled to the original sampling rate of EEG data (2,048 Hz) such that the HRV time course was synchronized with that of the EEG epochs. The HRV time series before and after upsampling were compared visually to confirm that no erroneous noise was introduced, as shown in S1 Figure. HR and HRV epochs from 1 s before to 20 s after cue onset were extracted. We set the following rejection criteria to ensure the quality of the HR and HRV indices. For each participant, two types of outliers were detected across and within trials and discarded according to robust z score criteria (57, 58): (1) *epochs* with absolute robust z scores larger than 3.4807 (equivalent to *p* = .0005) after applying a robust z transformation to the mean values of all epochs, and (2) epochs with robust z scores of any *sample* larger than the given threshold after applying a robust z transformation to the time series within the same epoch. The first step aimed to remove noisy sample points and maintain stationarity, while the second step aimed to remove outlier epochs. Finally, 14.42% of HR epochs and 19.64% of HRV epochs were discarded. A total of 205.12±13.51 HR epochs and 192.58±8.37 HRV epochs per participant were retained (see S1 Table and S2 Table for more details). These cleaned data were employed in subsequent analyses. The remaining HR and HRV epochs were downsampled to 200 Hz and averaged across epochs of the same condition within participants for subsequent analysis.

### 2.6. HEP computation

EEG signals were preprocessed using EEGLAB toolbox v2019.1 for MATLAB (59). Continuous raw data were bandpass-filtered (1–40 Hz), downsampled to 256 Hz, epoched relative to the cue onset (-1 s to 10 s), and baseline corrected (-1 s to 0 s) (Fig 2a). Epochs containing artifacts were detected by the automatic rejection algorithm implemented in the EEGLAB toolbox. Eye-related artifacts were corrected using automatic EOG correction with conventional recursive least squares regression. An extended Infomax independent component analysis was performed to remove EOG- and ECG-related components. Noise-free epochs were then rereferenced to the average of the EEG signals (Fig 2b). Finally, 9.55% of EEG epochs were discarded, and 217±14.82 epochs per participant were retained.

**Fig 2.**
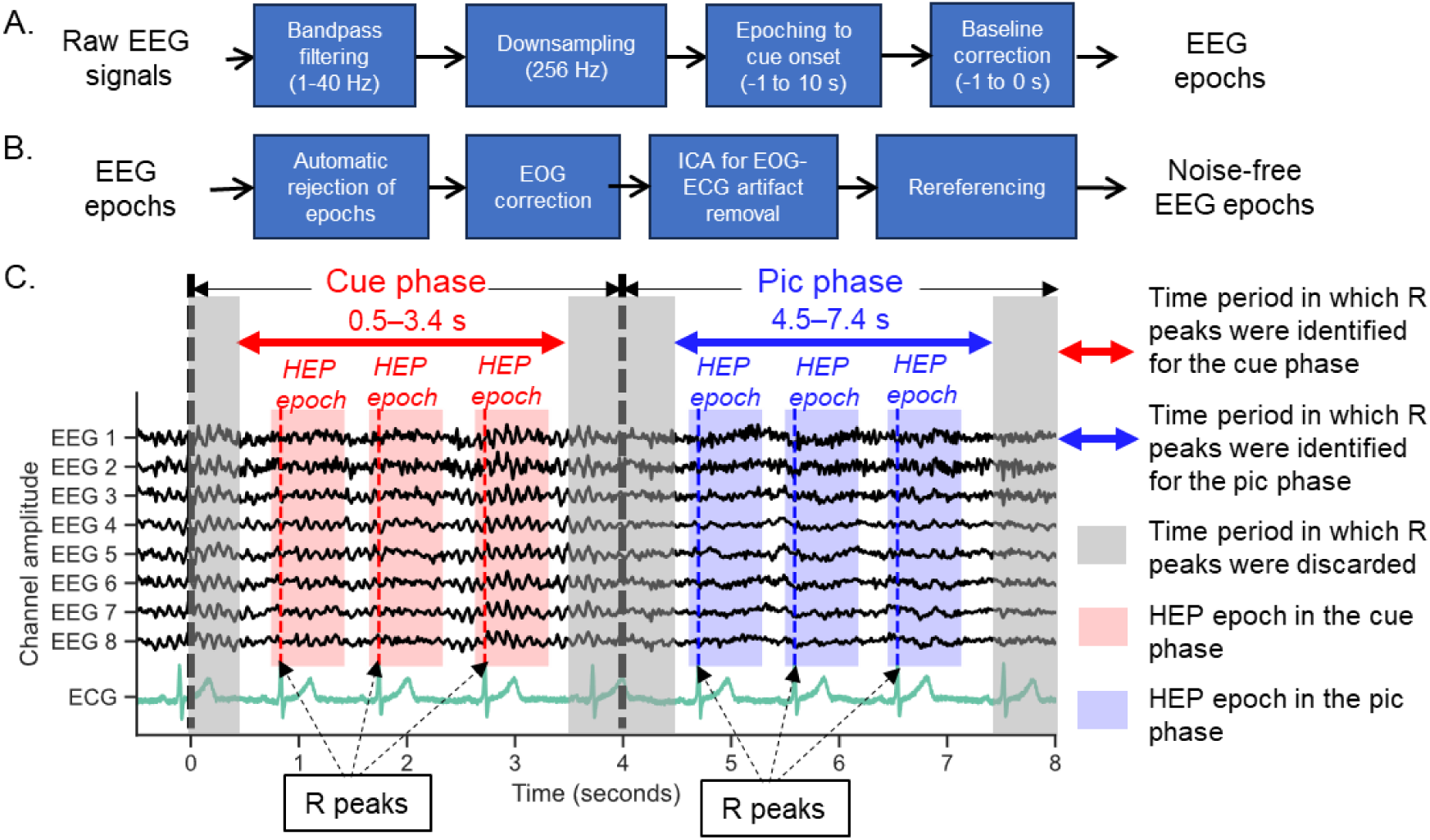
Flowchart of the processing of epochs to obtain heartbeat-evoked potentials (HEPs). (a) Preprocessing of raw EEG signals to obtain EEG epochs. (b) Artifact removal procedure used to obtain noise-free EEG epochs. (c) Procedures used to further epoch noise-free EEG epochs to obtain epochs containing HEPs within the cue presentation (Cue) phase and picture presentation (Pic) phase. For each of the EEG epochs, the time of each R peak was detected from the corresponding bandpass-filtered ECG signals for two periods: 0.5–3.4 s after cue onset (within the cue phase) and 4.5–7.4 s after cue onset (equivalent to 0.5–3.4 s after picture onset; during the pic phase) (Fig 2c). In this study, the R peaks with times close to the cue and picture onsets were not used in the HEP computation to minimize the direct perceptual effects of auditory and visual stimuli, respectively.

EEG-ECG epochs containing false-positive R peaks were identified and discarded based on careful visual inspection. On average, 2.46±9.50 EEG-ECG epochs were discarded per participant; 214.62±15.60 EEG-ECG epochs were retained. In the retained EEG-ECG epochs, the time points of R peaks were used as the onset time to further epoch the EEG signals (-0.1 to 0.6 s), followed by baseline correction (-0.1 to -0.05 s relative to the R peak onset). The baseline correction period was set at 0.05 s prior to R peaks to avoid cardiac artifacts. The HEPs were then computed separately for the corresponding conditions during the cue phase (PP_cue, PU_cue, UP_cue, and UU_cue) and pic phase (PP_pic, PU_pic, UP_pic, and UU_pic). The number of HEP epochs was 241.77±37.91 for each of the four predictable conditions (PP_cue, PP_pic, PU_cue, and PU_pic) and 120.71±19.86 for each of the four unpredictable conditions (UP_cue, UP_pic, UU_cue, and UU_pic). Note that onset times of HEP signals are the time points of the R peaks, as per convention, while those of cardiac reactivity values were the time points of cue onset.

To characterize the morphology of HEPs, real HEPs were compared with surrogate HEPs. Following the convention established by Gentsch, Sel (37), surrogate HEPs were computed based on surrogate R peaks that were randomly jittered around the real R peaks. In this study, the range of jittering was set to -0.5 to 0.5 s. Surrogate HEPs were epoched from the EEG-ECG epochs according to surrogate R peak times (S2 Figure). The averaged HEPs and surrogate HEPs were then obtained individually by averaging all real HEP epochs and surrogate HEP epochs, regardless of condition. A nonparametric cluster-level F test for spatiotemporal data implemented in MNE-Python (60) was then applied to the averaged real and surrogate HEP waveforms to identify clusters of channels and periods containing significant neural responses to heartbeats within the time window of 0.1 to 0.6 s after the R peak onset. Any electrodes located within 5 cm of each other were considered spatial neighbors. The number of permutations was set at 2,500, and the threshold to form spatiotemporal clusters was the F value corresponding to the significance threshold of *p* = .00001. After correction for multiple comparisons, the durations and sensor locations of the spatiotemporal clusters identified as significant at *p* < .05 were selected for further HEP analysis.

### 2.7. Statistical analysis

In this study, repeated-measures analyses of variance (RM-ANOVAs) and correlation and mediation analyses were performed using the Python package ‘pingouin’ (61). *t* tests and false discovery rate (FDR) corrections were implemented using the Python packages SciPy (62) and statsmodels (63), respectively. The α-level for FDR-corrected *p* values (*p*_FDR_) was set at 0.05. As needed, uncorrected *p* values (*p*_uncorrected_) less than 0.05 are also reported to describe trends, for completeness. The correlation analyses used in this study were partial correlation analyses controlling for age and sex, which have been shown to affect emotional processing (64–68), HEPs (69), and cardiac activity (70). The effect sizes of partial correlations were measured using *r*^2^.

#### 2.7.1. Assessment of whether cardiac reactivity is modulated by depression risk

To examine whether depression risk altered cardiac reactivity in response to the cues and emotional pictures, the interrelationship between cardiac reactivity and BDI scores was computed for each condition (PP, PU, UP, and UU). First, periods showing significant main effects of cues or emotional pictures were identified by performing RM-ANOVAs on the HR or HRV for each time point, followed by FDR correction for multiple comparisons. Since the recovery duration (the period until the waveform returned to baseline) for HR and HRV appeared to differ, we considered different time windows for HR and HRV: 12 s for HR waveforms and 20 s for HRV waveforms. For each of the significant periods identified, the correlation between each condition’s averaged cardiac reactivity and BDI scores was computed, followed by FDR correction for multiple comparisons.

#### 2.7.2. Relationships among depression risk, cardiac reactivity, and HEPs

For conditions that showed significant effects of depression risk on cardiac function, the interrelationship between depression risk and the activity of the cardiac interoceptive system was further quantified. The correlation between BDI scores and mean HEP amplitudes of each spatiotemporal HEP cluster was computed at each cue and pic phase. FDR correction for multiple comparisons was applied across clusters and phases.

To examine whether interactions between the cardiac interoceptive system and autonomic cardiac regulation were influenced by depression risk, mediation analysis was applied to examine whether the effect of HEPs on cardiac reactivity was mediated by depression risk, after controlling for age and sex. The analysis included the conditions, durations, and clusters showing significant correlations with BDI scores.

### 2.8. Control analyses

Cardiac activity affects the amplitudes of EEG signals and leads to cardiac field artifacts (CFA). Although careful preprocessing of EEG signals was performed to reduce CFA, we additionally analyzed mean ECG amplitudes to address the fact that ECG did not account for the differences in HEPs. We obtained ECG epochs from the bandpass-filtered ECG signals segmented using the time points of the R peaks as the onset times (-0.1 to 0.6 s), followed by baseline correction (-0.1 to -0.05 s). To test the effect of cues and emotional pictures on ECG amplitudes, RM-ANOVA was performed with post hoc tests to compare each time point during the 0.1–0.6 s interval, followed by FDR correction for multiple comparisons. RM-ANOVAs were also performed on ECG amplitudes of the R peaks, separately for the cue and pic phases. Moreover, correlations between BDI scores and the temporally averaged ECG amplitudes were computed for each time window corresponding to spatiotemporal HEP clusters. The correlations between BDI scores and R peak amplitudes and between BDI scores were also examined for each condition.

### 2.9. Correlation between HR and HRV

Previous studies have shown that HR and HRV are negatively correlated (71, 72). Moreover, interactions between cardiac activity measures vary according to tasks or subject groups (73). Most previous HR and HRV studies have computed average values across all epochs. However, our analysis was performed on the HR and HRV time series to identify which time windows showed significant effects. To examine the feasibility of the identified time windows, the correlation between the temporally averaged HR and HRV signals was computed for each condition; we expected to observe a negative correlation.

## 3. Results

### 3.1. Effect of cues, emotional pictures, and depression risk on HR

In all four conditions, HR slightly increased after cue onset and then decreased (Fig 3a). HR time series were significantly influenced by emotional pictures during 5.4–10.7 s after cue presentation (*p*_FDR_ < .05 and duration > 1 s for the main effect of pic in the cue × pic RM-ANOVA). During this period, unpleasant pictures elicited a larger deceleration of HR than pleasant pictures, especially in the unpredictable condition (UU > UP; *p*_FDR_ < .05 and duration > 1 s for the main effect of pic in the post hoc analysis; see Figs 3a and c). In the UU condition, the mean values of HR changes were negatively correlated with BDI scores (*r* = -.599, *p*_FDR_ = .008, *r*^2^ = .359, power = 0.920; see Fig 4a and Table 1). The high BDI group showed a larger deceleration in HR than the low BDI group (Fig 4c). In the other three conditions (PP, PU and UP), no significant correlation between HR changes and BDI scores was observed (Table 1). In the cue × pic RM-ANOVA of the HR time series, no main effect of cue was observed (*p*_FDR_ >.05).

**Fig 3.**
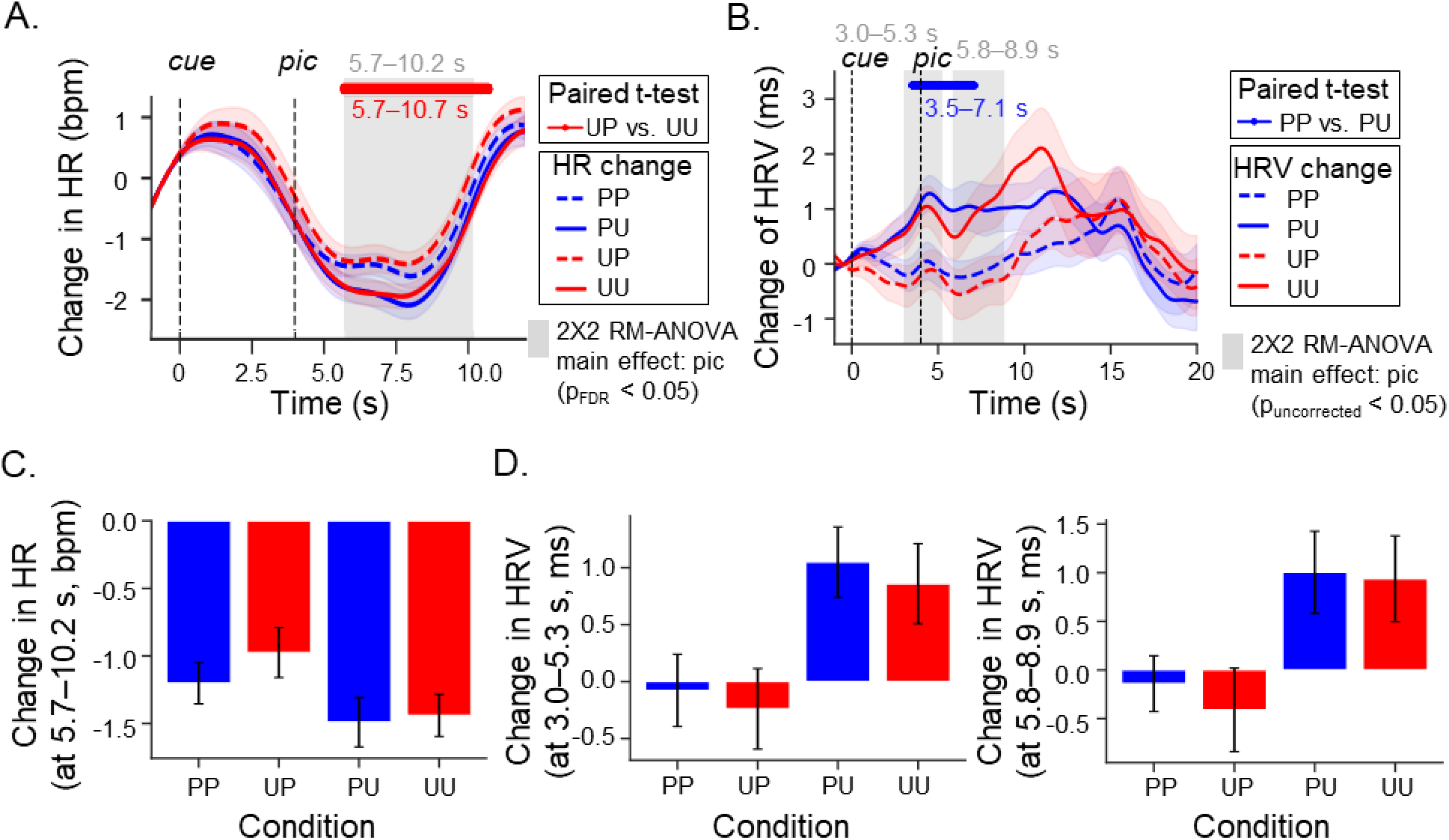
Changes of heart rate (HR) and heart rate variability (HRV). In the time series of (a) HR and (b) HRV, three periods show a significant main effect of picture valence in the cue × pic RM-ANOVA, as indicated by gray-shaded areas. The horizontal lines represent the period showing a significant difference according to paired *t* tests (*p* < .05 and duration > 1 s). The p values were FDR-corrected across time in the HR analysis. The p values were not corrected in the HRV analysis since none were significant after FDR correction. The left and right vertical dashed lines indicate the cue and picture onsets, respectively. Red and blue shaded areas along the HR and HRV time series represent one standard error from the means. The box plots represent temporally averaged (c) HR and (d) HRV values during periods showing significant main effects. PU and UU trials showed greater reductions in HR and increases in HRV than PP and UP trials. Red and blue colors represent predictable and unpredictable conditions, respectively. PP: predictably pleasant; PU: predictably unpleasant; UP: unpredictably pleasant; and UU: unpredictably unpleasant.

**Fig 4.**
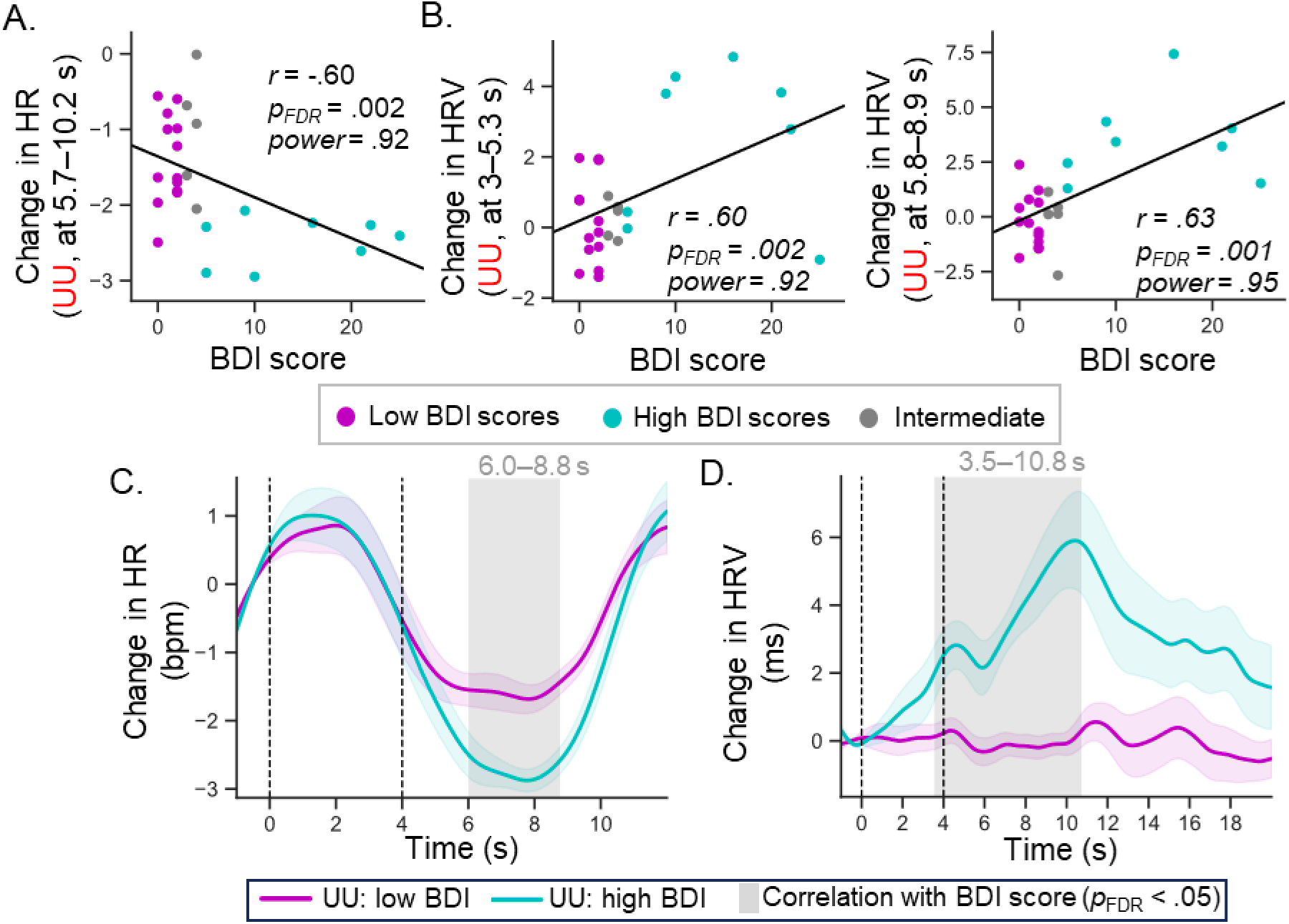
Relationships between BDI and changes in heart rate (HR) and heart rate variability (HRV). In the scatter plots of BDI scores with (a) heart rate (HR) changes or (b) heart rate variability (HRV) changes under the UU condition, the periods of HR and HRV were selected based on when the main effect of picture valence was significant in the cue × pic RM-ANOVA. The black solid lines represent fit lines for all samples. Cyan and magenta colors correspond to the high BDI group and low BDI group, respectively. The gray color indicates participants with intermediate BDI scores (between those of the two groups). In the time series showing (c) HR changes and (d) HRV changes under the UU condition in the low- and high-BDI groups, cyan- and magenta-shaded areas along the HR and HRV time series represent one standard error from the means. The gray-shaded area represents the duration showing significant correlations (FDR-corrected *p* < .05) of HR and HRV changes under the UU condition with BDI scores. BDI: Beck Depression Inventory; UU: unpredictably unpleasant.

**Table 1.** Correlation of BDI scores with mean HR changes during the 5.7–10.2 s interval, i.e., the period showing a significant main effect of picture valence in the RM-ANOVA, and HR changes in the low-BDI and high-BDI groups. *r* values are partial correlation coefficients after controlling for age and sex.

| Condition | Correlation with BDI scores |  |  |  |  | Change in HR (bpm) |  |
| --- | --- | --- | --- | --- | --- | --- | --- |
|  | *r* | *p* | *p*<sub>FDR</sub> | *r*<sup>2</sup> | Power | Low BDI group | High BDI group |
| PP | -.170 | .426 | .426 | .029 | .133 | -1.42 (1.02) | -1.50 (0.39) |
| UP | -.210 | .325 | .426 | .044 | .179 | -0.99 (0.79) | -1.55 (1.02) |
| PU | -.223 | .294 | .426 | .050 | .197 | -1.65 (1.04) | -2.42 (0.76) |
| UU | -.599 | .002<sup>+</sup> | .008* | .359 | .920 | -1.40 (0.59) | -2.46 (0.32) |

<sup>+</sup>: uncorrected $p < .05$ ; \*: FDR-corrected $p$ ( $p_{\text{FDR}} < .05$ ; data shown are the mean (*SD*).

### 3.2. Effect of cues, emotional pictures, and depression risk on HRV

HRV increased after cue onset in the unpleasant condition, but increased at approximately 10 s after cue onset in the pleasant condition (Fig 3b). The HRV time series were influenced by emotional pictures at 3.0–5.3 s and 5.8–8.9 s (*p*_uncorrected_ < .05 and duration > 1 s in the cue × pic RM-ANOVA for the main effect of pic). During these periods, unpleasant pictures elicited a larger increase in HRV than pleasant pictures, especially in the predictable condition (PU > PP; *p*_uncorrected_ < .05 and duration > 1 s in the post hoc analysis for the main effect of pic; see Figs 3b and d). During these periods, the mean HRV changes in the UU condition were significantly correlated with BDI scores (Table 2 and Table 3). Compared to the low BDI group, the high BDI group exhibited a larger increase in HRV in the UU conditions (Figs 4b and d). In the cue × pic RM-ANOVA of the HRV time series, no main effect of cue was observed (*p*_uncorrected_ >.05).

**Table 2.**
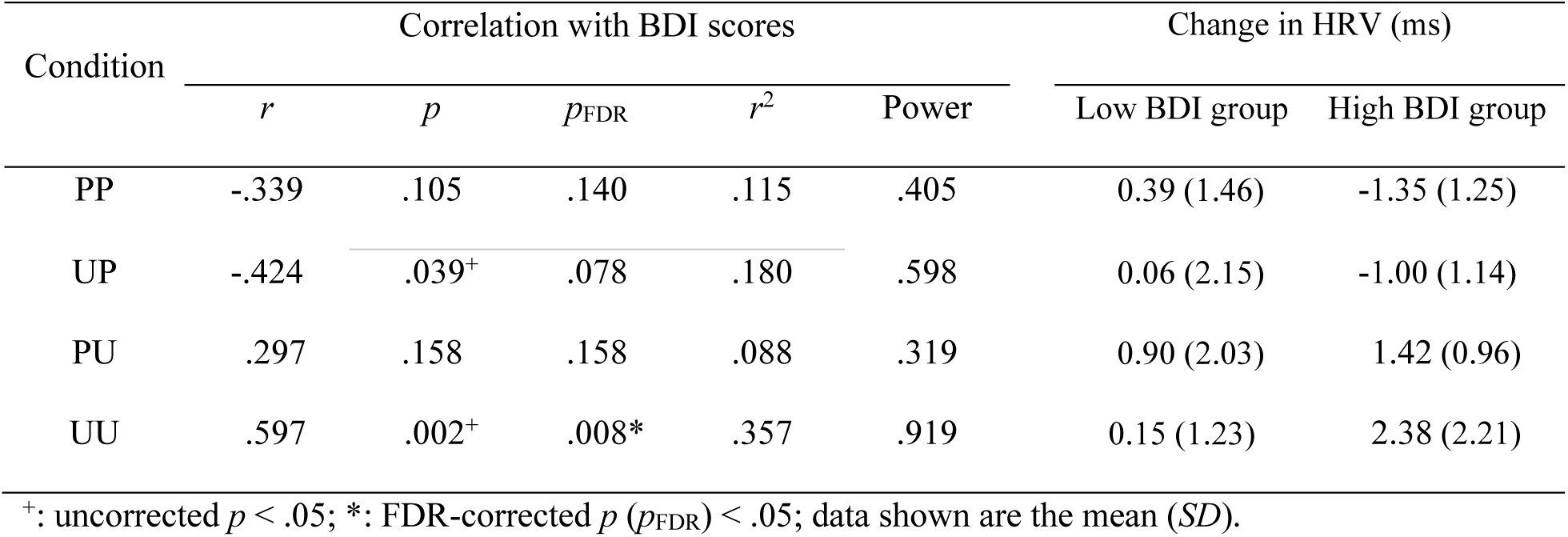
Correlation of BDI scores with mean HRV changes during the 3–5.3 s interval, i.e., the period showing a significant main effect of picture valence in the RM-ANOVA, and HRV changes in the low-BDI and high-BDI groups. *r* values are partial correlation coefficients after controlling for age and sex.

**Table 3.** Correlation of BDI scores with mean HRV changes during the 5.8–8.9 s interval, i.e., the period showing a significant main effect of picture valence in the RM-ANOVA, and HRV changes in the low-BDI and high-BDI groups. *r* values are partial correlation coefficients after controlling for age and sex.

| Condition | Correlation with BDI scores |  |  |  |  | Change in HRV (ms) |  |
| --- | --- | --- | --- | --- | --- | --- | --- |
| | $r$ | $p$ | $p_{\text{FDR}}$ | $r^2$ | Power | Low BDI group | High BDI group |
| PP | -.192 | .370 | .370 | .037 | .157 | 0.23 (1.03) | -0.96 (1.75) |
| UP | -.397 | .055 | .073 | .158 | .535 | 0.02 (1.93) | -1.67 (2.69) |
| PU | .511 | .011<sup>+</sup> | .021* | .261 | .785 | 0.22 (1.79) | 2.00 (2.69) |
| UU | .634 | .001<sup>+</sup> | .003* | .402 | .954 | -0.19 (1.23) | 3.46 (1.94) |

<sup>+</sup>: uncorrected $p < .05$ ; \*: FDR-corrected $p$ ( $p_{\text{FDR}} < .05$ ; data shown are the mean ( $SD$ ).

### 3.3. Spatiotemporal clusters of HEPs

Comparison of the grand averages of the real and surrogate HEP time series revealed that the real HEPs were stronger than the surrogate HEPs (Fig 5a). In the nonparametric cluster-level test comparing spatiotemporal data from real and surrogate HEPs (*p* < .05, duration > 0.1 s, number of channels ≥ 3), three significant spatiotemporal clusters were observed over the following locations: (1) the occipital and left temporoparietal regions during 0.10–0.38 s, (2) the central region during 0.10–0.29 s, and (3) the right temporal regions during 0.29–0.33 s (Fig 5b). Cluster 1 showed a positive deflection, while Clusters 2 and 3 showed negative deflections.

**Fig 5.**
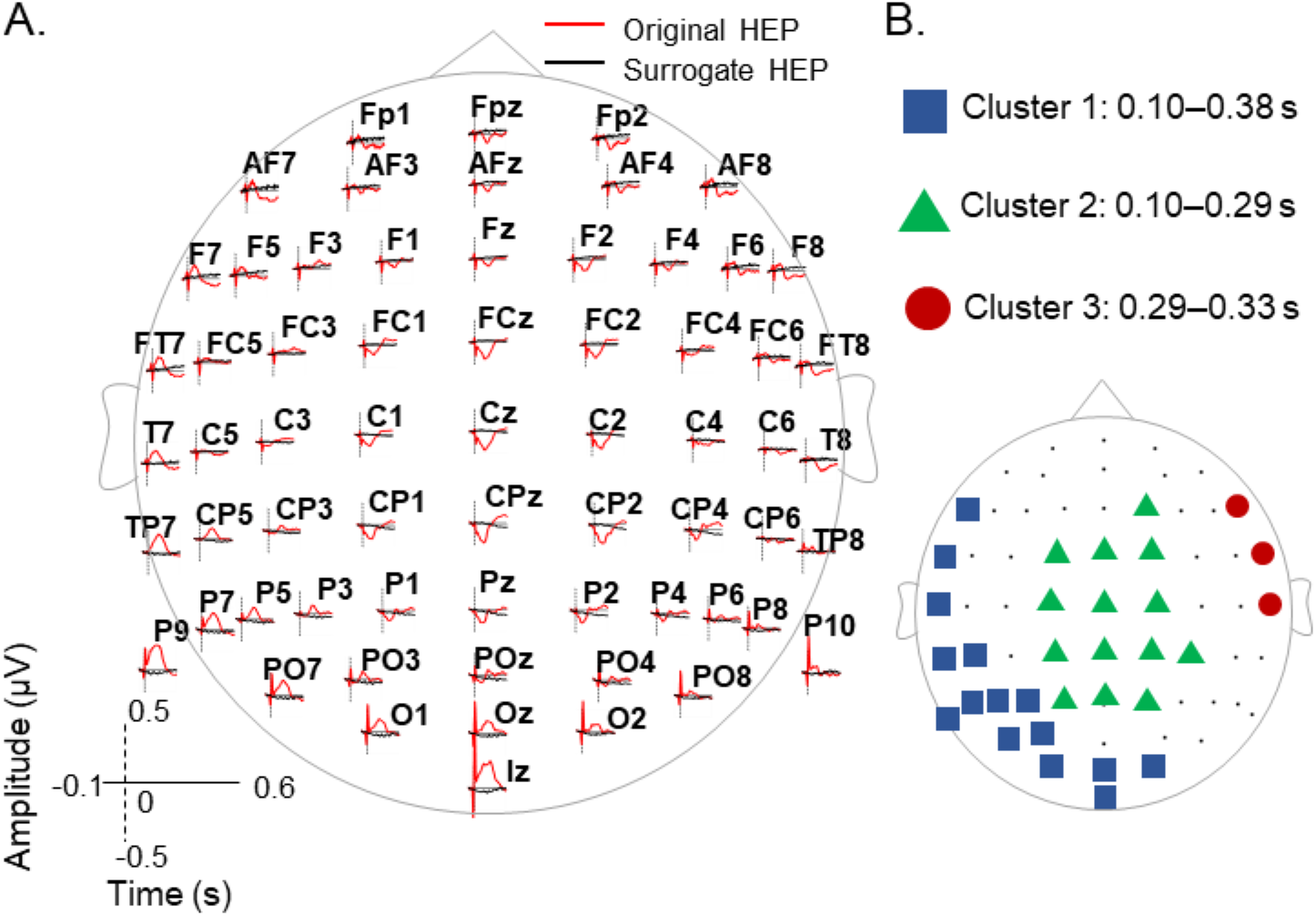
Time series of heartbeat-evoked potentials (HEPs) and channel locations of HEP clusters. (a) Grand average of original heartbeat-evoked potentials (HEPs), obtained by epoching EEG signals based on actual R peak times, and surrogate HEPs, obtained by epoching EEG signals based on randomly shifted R peaks, during the interval of -0.1 to 0.6 s. The scalp distribution of HEPs is shown from above. The text above each HEP plot indicates the name of the EEG channel. (b) Blue squares, green triangles, and red circles indicate channel positions of HEP clusters 1, 2, and 3, determined by the spatiotemporal nonparametric cluster-level test of the time series of HEPs and surrogate HEPs during 0.1–0.6 s. Significant durations for HEP clusters 1, 2, and 3 are 0.10–0.38 s, 0.10–0.29 s, and 0.29–0.33 s, respectively.

### 3.4. Effect of depression risk on HEP clusters

In Sections 3.1 and 3.2, we reported that the decrease in HR and increase in HRV in the UU condition were correlated with BDI scores. Focusing on the UU condition, we further examined the relationship between BDI scores and mean amplitudes of HEP clusters. We found that HEP cluster 1 was correlated with depression risk when processing nonpredictive cues (UU_cue, *r* = -.48, *p*_uncorrected_ = .017, *p*_FDR_ = .033; see also Fig 6a and Table 4). The positive HEP component increased to a lesser extent in the high BDI group than in the low BDI group. Since the amplitude of event-related potentials (ERPs) may be affected by the thickness of the skull, the HEP cluster 1 for the PU condition was additionally plotted as the baseline of that group for visual inspection. As shown in Fig 6b, the HEP amplitudes in the UU condition were smaller than those in the PU condition in the high BDI group. On the other hand, in the low BDI group, the HEP amplitudes showed a larger increase in the UU condition than in the PU condition. Overall, the HEP amplitudes of the left occipital-temporal-parietal region when processing nonpredictive cues decreased with increasing depression risk.

**Fig 6.**
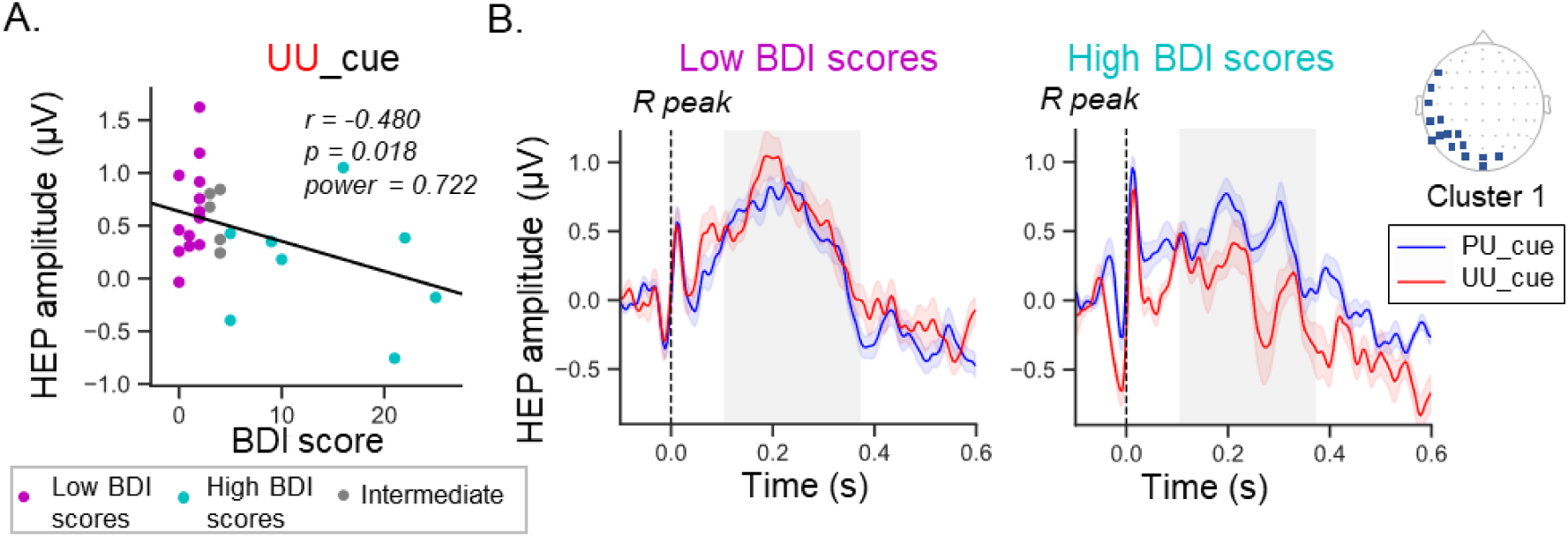
Relationships between BDI and heartbeat-evoked potentials (HEPs). (a) Scatter plot of BDI scores with the amplitude of heartbeat-evoked potentials (HEPs) in cluster 1 under the UU condition during the cue phase. Cyan and magenta colors indicate the high BDI group and low BDI group, respectively. Gray circles indicate participants with intermediate BDI scores, that is, scores between the thresholds of the high BDI group and low BDI group. (b) Time series of averaged HEPs across electrodes of cluster 1 in the PU and UU conditions during the cue phase. Gray-shaded areas indicate the duration of HEP cluster 1. Blue- and red-shaded areas along the HEP time series indicate one standard error from the mean. Dashed lines indicate the time of R peaks. BDI: Beck Depression Inventory.

**Table 4.**
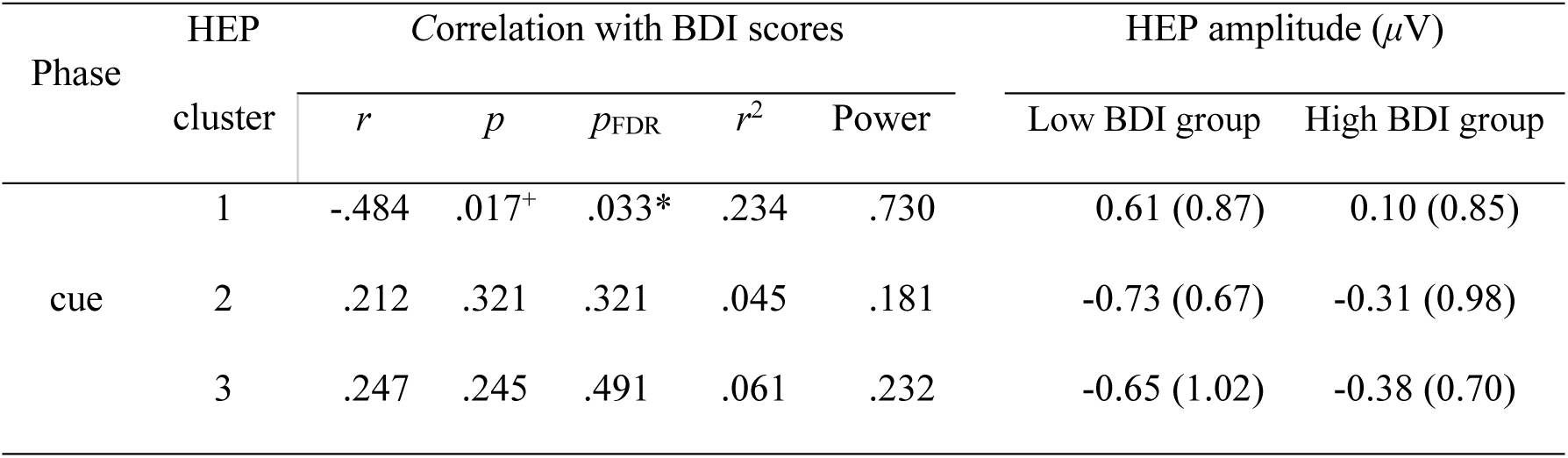
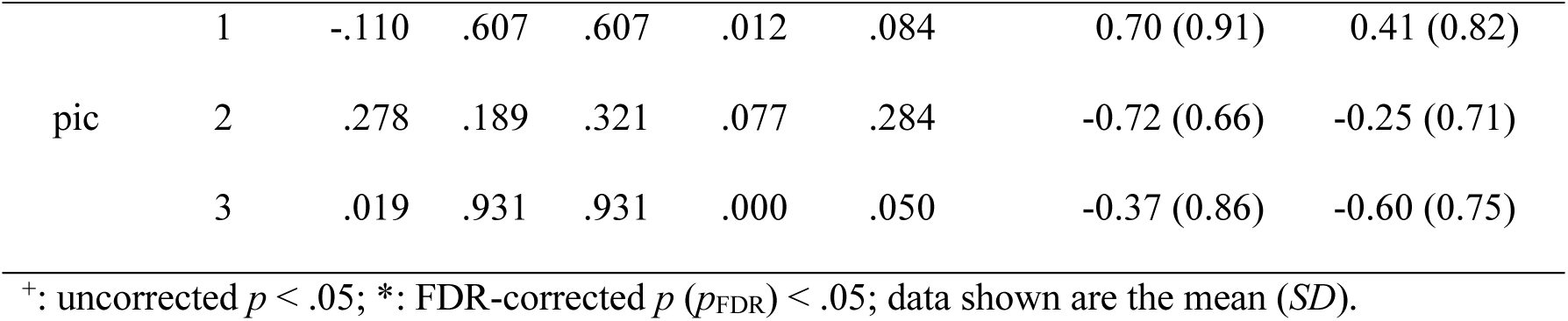
Correlation between BDI scores and mean HEP amplitudes in the *UU* condition and for each cluster during the *cue* and *pic* phases as well as the HEP amplitudes in the low-BDI and high-BDI groups. *r* values are partial correlation coefficients after controlling for age and sex.

### 3.5. Depression risk mediated the relationship between HEPs and cardiac reactivity

In the UU condition, the mean HEP amplitudes during the cue phase, the mean HR changes, and the mean HRV changes tended to predict BDI scores (Figs 4a–b and 6a). Thus, we further examined the mediating effect of BDI scores on the relationship between HEPs in cluster 1 and cardiac reactivity in the UU condition. This analysis showed that the indirect effect of mean HEP amplitudes on HR changes was significantly mediated by depression risk, whereas the direct effect was nonsignificant (Fig 7a). Moreover, the interaction between mean HEP amplitudes and mean HRV changes was also mediated by depression risk, and the direct effect was also nonsignificant (Fig 7b). Since cardiac reactivity may also influence HEPs, we additionally assessed the mediating effect of BDI scores on the relationship between cardiac reactivity and HEPs during the picture phase. In this analysis, no significant indirect effect was found (S3 Figure). Overall, our results suggest that BDI scores mediate the association between HEPs during the prediction phase and cardiac reactivity in response to unexpected, unpleasant pictures.

**Fig 7.**
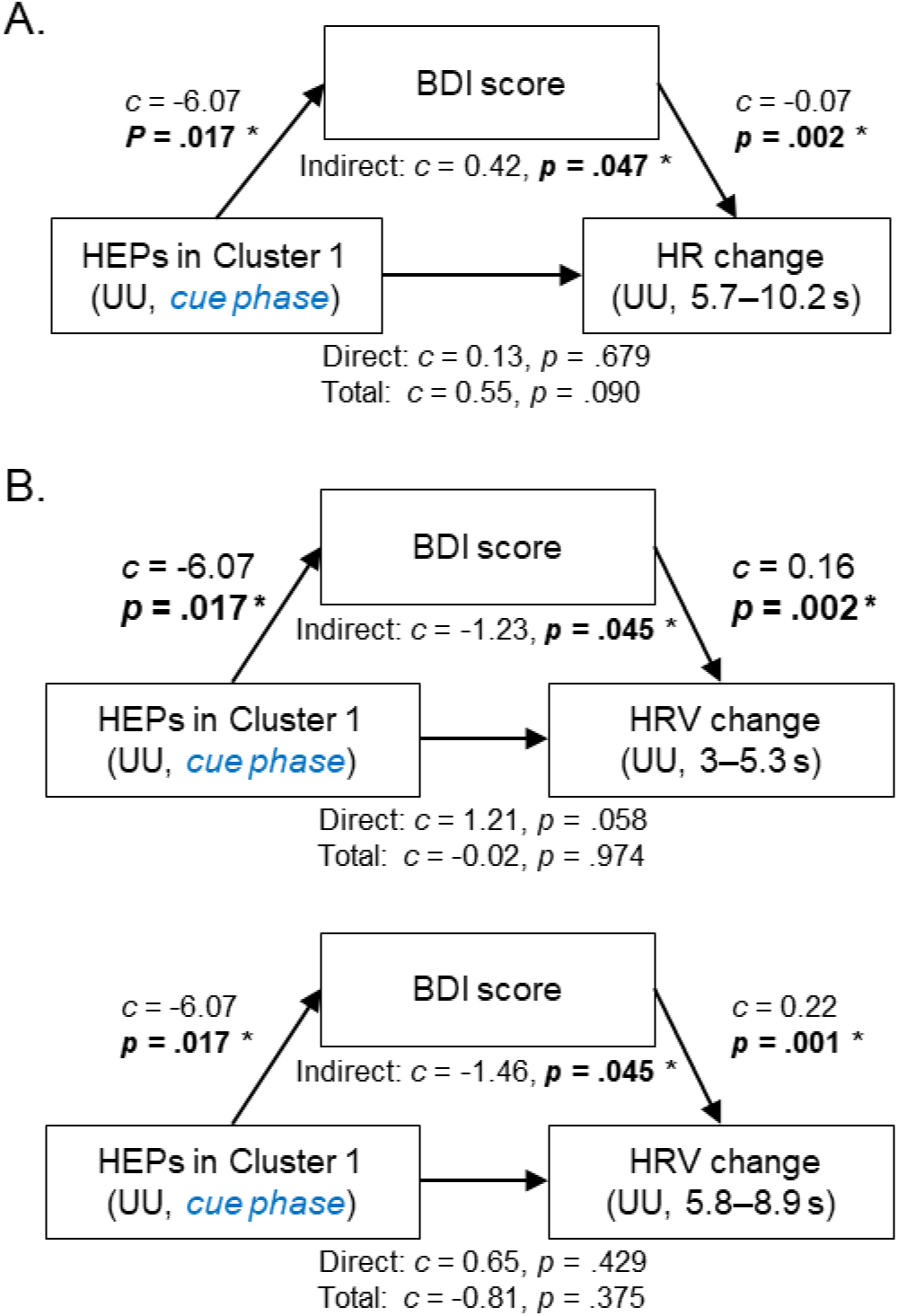
Results of mediation analysis. Mediation models predicting changes in (a) heart rate (HR) and (b) heart rate variability (HRV) under the unpredictably unpleasant (UU) condition from the heartbeat-evoked potential (HEPs) in cluster 1 at the *cue phase* mediated by depression risk, which was quantified by the BDI score. *: *p* < .05. BDI: Beck Depression Inventory.

### 3.6. Effects of cues, emotional pictures, and depression risk on ECG signals

The RM-ANOVA including the factors of cue and the emotional valence of pictures did not reveal any significant main effects on ECG data at any time point within the period of 0.1–0.6 s (S4 Figure). The paired *t* tests on the ECG time series also did not reveal significant differences between predictable and unpredictable conditions. Moreover, when RM-ANOVA was performed with the factors of cue and the emotional valence of pictures on the ECG amplitudes of R peaks, the main effects of cue and picture valence, as well as their interaction, were not significant in either the cue or pic phases (S3 Table). The ECG amplitudes at R peaks were also not significantly correlated with BDI scores in any of the conditions (S4 Table). Last, in the condition in which BDI scores were found to modulate the effects on HEPs, i.e., the UU condition, we examined the correlation of BDI scores and ECG amplitudes averaged over the periods in which spatiotemporal HEP clusters were identified, and none of the correlations were significant (S5 Table). Thus, the results of the control analysis suggest that the modulating effects on HEP amplitudes were not driven by ECG artifacts and may instead represent modulations of cortical processing of heartbeat signals.

### 3.7. Nonsignificant negative correlations between HR and HRV

As expected, the mean magnitudes of HR changes were negatively correlated with HRV changes during the time windows in which a significant main effect of picture valence was identified by RM-ANOVAs (HR: 5.7–10.2 s; HRV: 3–5.3 s and 5.8–8.9 s; see S5 Figure). Thus, the relationships between HR and HRV changes in our selected time windows aligned with those reported in previous studies, suggesting that our data-driven approach for selecting time windows for further analysis was feasible.

## 4. Discussion

This study examined the extent to which depression risk is associated with cardiac and neurophysiological emotional responses in certain and uncertain circumstances. A comprehensive temporal characterization of HR and HRV responses revealed dissociable and unique relationships with the valence of emotional pictures. HDR was associated with a larger reduction in HR, a larger increase in HRV and a smaller HEP amplitude in response to unpredictable, unpleasant stimuli. This finding supports the hypothesis that depression risk alters interoceptive processing and increases physiological fluctuations in response to unpredictable stimuli. Depression risk appeared to influence anticipatory HEP responses and mediate the relationship between the anticipatory HEP and poststimulus cardiac reactivity, especially in response to unexpected, unpleasant pictures (i.e., in UU trials). Importantly, the abovementioned HEP findings were not explained by any ECG features, confirming the unique role of HEPs. These results suggest that individuals with HDR have reduced neural resources for anticipation and increased cardiac reactivity following unpredictable negative events. The present study provides evidence that disrupted cardiac regulation and interoception are indicators of HDR.

### 4.1. Depression risk modulated neurophysiological reactivity to negative events

In line with our expectations, modulating effects of BDI scores on HR and HRV were observed in negative emotional conditions but not in positive emotional conditions. A correlation of the intervals between R peaks and self-reported valence was reported during the processing of negatively valenced stimuli but not positively valenced stimuli (74), indicating stronger modulation of cardiac activity during the generation of negative emotion. Thus, the processing of negatively valenced stimuli may rely on interoceptive information more than the processing of positively valenced stimuli. Our mediation analysis results suggested that prestimulus cardiac interoceptive processing affected poststimulus cardiac activity in response to negatively valenced stimuli and further showed that the interaction was mediated by BDI scores. Attentional bias toward negative events is a notable characteristic of individuals with MDD (75) or subclinical depression (76). Therefore, our findings suggest that individuals with high BDI scores exhibited altered autonomic cardiac functions and cardiac interoception, as reflected by neural and physiological cardiac responses to unpleasant stimuli.

Altered HEPs and cardiac reactivity to unpredictable negative events may be particularly relevant when modeling the relationships of interoceptive system dysfunction and atypical autonomic regulation with the risk of depression. Abnormal cardiac activity (32, 77, 78) and low interoceptive sensitivity (79–81) have been widely documented in depressed patients, suggesting associations of depression with autonomic imbalance and interoceptive impairment. Our findings enhance understanding of this relationship, revealing that disturbances in autonomic cardiac functioning and the brain interoceptive network were also observed in healthy individuals with high BDI scores. Although the causal relationships of depression with ANS dysfunction and interoceptive system impairment have not been confirmed, some longitudinal evidence has suggested that dysfunction of the ANS and interoceptive system may precede the onset of depression. Recently, two longitudinal studies showed that resting-state HRV predicted subsequent depressive symptoms (78, 82), suggesting that disruption of the ANS may increase susceptibility to depression. Somatic symptoms, which have been related to dysfunction of the interoceptive system, can also predict subsequent depression (83, 84), implying that there is a link between abnormal interoceptive system function and HDR. The exact mechanisms by which disturbances in the ANS and interoceptive system may increase depression risk are still unknown. Disruptions of the ANS can induce inflammation (85) and metabolic dysregulation (86). Low sensitivity to interoceptive signals may lead to disrupted predictive coding of interoceptive signals (19, 87), which is crucial to emotional processing and allostatic regulation.

Moreover, our mediation analysis results showed that the association between HEPs and cardiac reactivity was mediated by BDI scores under uncertain conditions (Fig 7), suggesting that brain–heart communication was influenced by depression risk. This may result from the enhanced neural control of cardiac activity as depression risk increases, which has been found in patients with subclinical depression during the resting state (88). Moreover, reduced baroreflex sensitivity (BRS) has been reported in depressed patients (89, 90). The baroreflex is one of the key mechanisms to maintain homeostasis and reduce fluctuations in blood pressure (BP) (91). Decreased BP reduces the activation of baroreceptors and their effects on the central inhibitory pathway, leading to increased sympathetic tone, HR, and BP. Minati et al. (92) reported reduced BP and HR during the processing of negative emotions. Notably, the trend observed in the HR time series in our study is consistent with previous results (92). Moreover, BP responses were previously found to be correlated with baroreceptor activation of the central feedback pathway in emotional picture-perception tasks (93). Increased BP variability has been related to increased severity of subclinical depression, suggesting a link between baroreflex dysfunction and HDR (94). Thus, we speculated that individuals with high BDI scores also have reduced BRS. In individuals with low BDI scores, HR changes in response to negative stimuli may be partly attributed to the baroreflex. In contrast, in individuals with high BDI scores, a low BRS may reduce the effect of baroreceptors on the brain, leading to altered brain–heart communication. Baroreceptor dysfunction may also explain the exaggerated cardiac reactivity or low capacity to maintain homeostasis in these individuals. Further investigation of the relationship between the baroreflex and depressive symptoms are needed to identify the mechanisms underlying abnormal autonomic and interoceptive activity. Baroreflex sensitivity or BP should be measured in future studies.

### 4.2. Low anticipatory HEP amplitudes in individuals with HDR

The low anticipatory HEP amplitudes observed in individuals with high BDI scores may imply that they have a reduced sensitivity to cardiac afferent signals (Figs 6a–b). Notably, compared to healthy controls, MDD patients have exhibited reduced HEP amplitudes (31). In contrast, among healthy participants, HEP amplitudes were not altered with depression level (88, 95). The differences between studies may result from the use of different tasks, electrodes, and selected time windows. Catrambone et al. (88) compared HEP amplitudes averaged over 250 to 500 ms during the resting state, and Judah et al. (95) selected HEPs from electrode *Fz* averaged over 200 to 350 ms. The resting state has been associated with low interoceptive signal intensity, which increases the difficulty of identifying individual differences in cardiac interoception; thus, increasing sympathetic activity through tasks, such as the emotion-evoking picture-evaluation task employed in our study, may facilitate cardiac interoception (96). Furthermore, as shown in Fig 5b, the HEP cluster negatively correlated with BDI scores in the UU condition was distributed in the occipital, left temporoparietal, and left frontolateral regions. HEP amplitudes at electrodes over left temporal and frontolateral regions have been suggested to reflect the afferent cortical representation of myocardial function (97).

The weak anticipatory HEP amplitudes observed in individuals with high BDI scores may result from reduced attention to cardiac signals and enhanced attention to incoming external events. HEP amplitudes appeared to be enhanced when participants attended to their heartbeats rather than to external stimuli (98). The HEP amplitude is considered a neural correlate of predictive processing (99, 100). In the model proposed by Barrett et al. (19), depression was associated with inefficiency of the prediction model, due to low precision of predictive signals and/or deviations in the prior (101, 102). Precision, i.e., the inverse of uncertainty, is built up by environmental reliability and modulated by attention (26, 98, 103). Depressive traits may correspond with *reduced* precision of predictions and *increased* precision of prediction errors (102). Therefore, our results imply that HDR is associated with reduced precision or defective priors regarding interoceptive prediction, which may emerge from long-term discrepancies between predictions and prediction error in individuals with HDR (102). Researchers have reported that individuals with HDR may exhibit altered anticipatory responses in the insula (INS), ventrolateral prefrontal cortex, anterior cingulate cortex, and ventral striatum (39, 40). The INS is considered one of the major neural sources of the HEP (104) and is responsible for the prediction and prediction error processing of current and future physiological status (36, 105). Thus, our findings provide additional neurophysiological evidence that alterations in cardiac interoception are associated with depression risk. Further investigation of the cortical sources of the left temporoparietal HEP may clarify whether associations between anticipatory HEPs and depression risk depend on the interoceptive brain region.

### 4.3. Effects of valence on the HR and HRV time series

Our post hoc analysis of the HR time series confirmed earlier findings that unpleasant (negatively valenced) pictures *decreased* HR more than pleasant (positively valenced) pictures (22, 106–108). Some studies have reported a larger deceleration of HR in response to unpredictable stimuli compared to predictable stimuli (35, 36). However, we did not observe a difference in HR deceleration between the unpredictable and predictable conditions (Fig 3a). This discrepancy with previous studies may have resulted from the use of different stimuli. For example, participants in another study were presented with anticipated painful thermal stimuli (36), which may induce higher arousal and stronger negative emotions than negative picture stimuli. Another possible reason is the difference in the selected baseline periods. Some previous studies have used the period before the presentation of the pictures as a baseline. Since anticipation induces changes in physiological activity (109, 110), our design may be more sensitive to the effects of anticipation. Future research on physiological activity should consider the effects of anticipation when a variety of cue types, each representing different levels of certainty or upcoming stimuli, are used.

Our analysis of the HRV time series yielded results inconsistent with those of previous studies, showing that the HRV in response to viewing positively valenced images was higher than that in response to viewing negatively valenced images (111). Moreover, another study reported reduced HRV after viewing emotional pictures (25). These differences may be caused by differences in signal processing methods and paradigms. Previous studies have computed the averaged HRV over a long trial. Instead, we computed HRV time series by applying a 10-s sliding window, which could reveal the temporal dynamics of cardiac reactivity. Moreover, in previous experiments, (25) and (111), positive or negative affect was induced via exposure to 5 min excerpts of a speech or video. Thus, cardiac regulation may show different long-term and short-term changes or different responses to mood and emotion.

### 4.4. Altered predictions and prediction error processing in individuals with HDR

Furthermore, our correlation analysis showed that the HRV changes in the UU condition increased as the BDI score increased (Fig 4b), suggesting that prediction error signals elicited by experiencing unexpected negative emotions are altered by depression risk. Additionally, the correlation between the HEP amplitude during the anticipation period and BDI scores (Fig 6a) may indicate that the neural processing of prediction is modulated by depression risk. Moreover, the relationship between the neural signals comprising predictions and cardiac signals comprising prediction errors was mediated by BDI scores (Fig 7b). Future studies using computational models (112, 113) may be needed to examine alterations of predictive coding associated with depression risk based on neuro-cardiac interactions.

### 4.5. Imbalance of parasympathetic and sympathetic nervous system activity in individuals with HDR

We also examined temporal characteristics of HR and HRV changes during anticipation according to predictive cues and perception of emotional pictures. The effects of anticipation on HR appeared soon after the presentation of cues, and differences in HR deceleration according to the emotional pictures (pleasant vs. unpleasant) appeared at approximately 5.7 s, followed by the recovery of HR after 10–12 s (Fig 3a).

Cardiac activity is remotely controlled by the central autonomic network via the regulation of the ANS (114). The two branches of the ANS, the parasympathetic (PNS) and sympathetic nervous systems (SNS), act antagonistically to regulate cardiac activity. It is well known that activation of the PNS decreases HR, whereas activation of the SNS increases HR. The PNS alters cardiac signals after 1 s, while the effect of the SNS is slower, altering cardiac signals after 5 s (115). Thus, the HR deceleration initiated during the cue phase may have been modulated mainly by the PNS activation, and the recovery of HR may reflect the activation of the SNS. HRV increased after cue onset and peaked after approximately 10 s in the UU condition (Fig 3b). SDNN, the measure of HRV used in this study, is thought to reflect both SNS and PNS activation (115). Overall, the effects of the PNS and SNS were reduced approximately 10 s after cue onset or approximately 6 s after the onset of the unexpected, unpleasant pictures. Moreover, the larger reduction in HR and stronger increase in SDNN in individuals with high BDI scores may indicate heightened PNS activity elicited by exposure to unexpected negative stimuli among HDR individuals.

### 4.6. Limitations

Despite the comprehensive nature of our analyses, this study has several limitations. First, the standard method used to calculate short-term HRV from 5-min ECG recordings (53) was not feasible in our event-related paradigm. However, it has been suggested that accurate measures of SDNN can be obtained from 10-s ECG recordings (55, 56); thus, we computed the temporal dynamics of HRV using a 10-s sliding window. Our analysis of HRV from ultrashort (10-s) ECG recording segments may not be directly comparable to the standard short-term HRV measures obtained from 5-min ECG recordings. However, ultrashort HRV values have been increasingly applied in investigations of cardiovascular diseases (116–118). Second, our analysis of cardiac reactivity was limited to two time-domain parameters, HR and SDNN. Other cardiac reactivity markers for HRV, such as frequency-domain parameters (e.g., low- or high-frequency components or nonlinear geometric methods (115)), might have been used. However, these measures typically require ECG recordings of at least 60 s (55), so they would not fit well with our experimental design. Therefore, future studies could assess these measures using different experimental designs, such as a blocked design with a longer intertrial interval, rather than the event-related design we employed here. Third, although we tried to dissociate the event-related effects of the cues and pictures on cardiac reactivity, temporal dissociation of these two events might not have been completely achievable because of the <u>fixed</u> time interval between cues and pictures in our paradigm.

Fourth, potential comorbid conditions of depression were not considered in this study. Psychological and physical factors, such as anxiety level and body mass index, may also influence the HEP effect we observed. Fifth, while we found associations of HEPs and cardiac responses with subjective depressive risk (assessed by BDI scores), such subjective ratings are subject to human bias; thus, future studies could investigate this relationship using other, more objective scoring systems, such as the Hamilton Rating Scale for Depression (HAM-D) (119). Sixth, the nonstationary nature of brain activity elicited by a randomized sequence of stimuli (120) was not investigated in this study. To minimize the effect of nonstationarity, the order of trials was randomly selected for different participants. Moreover, the event-related signals were processed using highpass filters and baseline correction to remove any long-term trends caused by nonstationarity, and waveforms were computed by averaging across multiple trials within the same condition. Further investigations are needed to disentangle the temporal characteristics of nonstationarity. Although this study had some limitations and discrepancies, our evidence suggests that HDR affects interoceptive cardiac processing through responses to cues and emotional pictures.

### 4.7. Conclusion

In conclusion, our findings revealed that HDR is associated with enhanced cardiac reactivity, reduced anticipatory HEP amplitudes, and altered neuro–cardiac interactions in response to unexpected, unpleasant stimuli. These alterations may correspond to hyperactivity of the PNS, a reduced BRS, and ineffective interoceptive predictive coding. Our findings suggest that alterations in the ANS and cardiac interoceptive system, as well as their interaction, may reflect depression risk. These indices of cardiac and neuro–cardiac reactivity to unexpected negative stimuli may be useful in neurophysiological feedback treatment for depression. Our findings enhance understanding of emotional processing under certain and uncertain conditions in the healthy population and revealed a novel link between depression risk and brain-heart interactions through detailed characterization of cardiac reactivity and HEP time series.

## Acknowledgments

This study was supported by the Japan Science and Technology Agency (JST) COI (grant nos. JPMJCE1311 and JPMJCA2208), JST Moonshot Research and Development Program (grant no. JPMJMS2296), KAKENHI Grant-in-Aid for Early-Career Scientists (grant no. 20K16627), and in part by the Higher Education Sprout Project, funded by the Ministry of Education, at the Headquarters of University Advancement at National Cheng Kung University (NCKU). We thank Kai Makita at the University of Fukui, Japan, for data collection. CHL participated in study design and data analysis and wrote the paper. NK participated in study design and data collection. RM participated in data analysis. SY obtained the research grants from the COI and Moonshot programs, supervised the research, and helped write the paper. MGM participated in study design and carefully revised the paper. The authors have no conflicts of interest to declare regarding this research.

## Supporting information captions

**S1 Table.** Numbers of epochs of heart rate (HR) time series retained and removed after outlier detection using robust z score criteria.

**S2 Table.** Numbers of epochs of heart rate variability (HRV) time series retained and removed after outlier detection using robust z score criteria.

**S3 Table.** Results of the cue × pic RM-ANOVA on ECG amplitudes at R peaks.

**S4 Table.** Correlation between BDI scores and ECG amplitudes at R peaks for the UU, PU, UP, and UU conditions at the cue and pic phases. BDI: Beck Depression Inventory; PP: predictably pleasant; PU: predictably unpleasant; UP: unpredictably pleasant; and UU: unpredictably unpleasant.

**S5 Table.** Correlation between BDI scores and ECG amplitudes averaged over periods for HEP clusters in the UU condition at the cue and pic phases. BDI: Beck Depression Inventory; UU: unpredictably unpleasant.

**S1 Figure. Comparison of the heart rate variability (HRV) time series before and after upsampling**. HRV time series were upsampled from 4 Hz to 2,048 Hz over (a) a short segment (5 s) and (b) a long segment (2,500 s). The data are from one of the participants. The top and bottom panels show the time series before and after the upsampling process, respectively. In (a), each square represents a single sample of HRV. The squares in the bottom overlap with each other and thus resemble a thick line. These figures show that the HRV after upsampling was similar to that before upsampling.

**S2 Figure. Diagram showing how real and surrogate heartbeat-evoked potential (HEP) epochs were calculated.** HEP epochs were obtained by segmenting EEG epochs during the period of -0.1 to 0.6 s with the true R peak time as the onset time, followed by baseline correction. Surrogate R peaks were jittered from true R peaks in the range of *-*0.5 to 0.5 s (green dashed arrows). Surrogate HEP epochs were then obtained by segmenting EEG epochs during the interval of *-*0.1 to 0.6 s, with surrogate R peak times as the onset time, followed by baseline correction. Note that the R peaks and surrogate R peaks occurring outside the ranges of 0.5–3.4 s and 4.5–7.4 s were discarded.

**S3 Figure. Results of mediation analysis.** The analysis examining whether depression risk mediates the relationship between cardiac reactivity, reflected by (a) heart rate (HR) and (b) heart rate variability (HRV), and neural responses to emotional pictures, represented by heartbeat-evoked potentials (HEPs) in the picture (pic) phase. No significant mediation effect was found in these three models. BDI: Beck Depression Inventory; UU: unpredictably unpleasant.

**S4 Figure. Time series of averaged ECG epochs for the PP, PU, UP and UU conditions.** The vertical black dashed line indicates the time of the R peaks. PP: predictably pleasant; PU: predictably unpleasant; UP: unpredictably pleasant; and UU: unpredictably unpleasant.

**S5 Figure. Scatter plots of changes in heart rate (HR) and heart rate variability (HRV).** The changes in HR were averaged over the interval of 5.7–10.2 s and those in heart rate variability (HRV) were averaged over the intervals of (a) 3–5.3 s and (b) 5.8–8.9 s in the unpredictably unpleasant (UU) condition. Black solid lines represent regression fits of data from all participants.

