## Supporting information for "Brain–heart dynamics during emotional processing under uncertain conditions: An index of depression risk"

**S1 Table.** Numbers of epochs of heart rate (HR) time series retained and removed after outlier detection using robust z score criteria.

| Condition | *Number of retained epochs*  *(standard deviation)* | | |  | *Number of removed epochs*  *(percentage)* | |
| --- | --- | --- | --- | --- | --- | --- |
|  | *Original* | *Criterion 1* | *Criterion 2* |  | *Criterion 1* | *Criterion 2* |
| PP | 79.8 (0.4) | 79.1 (1.4) | 68.8 (4.9) |  | 0.8 (1.0%) | 10.3 (13.0%) |
| PU | 80.0 (0.0) | 79.2 (1.1) | 68.3 (5.3) |  | 0.8 (1.0%) | 11.0 (13.8%) |
| UP | 39.9 (0.3) | 39.5 (0.7) | 33.8 (3.0) |  | 0.4 (1.0%) | 5.7 (14.4%) |
| UU | 39.9 (0.3) | 39.6 (0.7) | 34.2 (2.9) |  | 0.3 (0.8%) | 5.4 (13.8%) |

**S2 Table.** Numbers of epochs of heart rate variability (HRV) time series retained and removed after outlier detection using robust z score criteria.

| Condition | *Number of retained epochs*  *(standard deviation)* | | |  | *Number of removed epochs*  *(percentage)* | |
| --- | --- | --- | --- | --- | --- | --- |
|  | *Original* | *Criterion 1* | *Criterion 2* |  | *Criterion 1* | *Criterion 2* |
| PP | 79.8 (0.4) | 78.8 (2.0) | 64.0 (4.3) |  | 1.0 (1.3%) | 14.8 (18.9%) |
| PU | 80.0 (0.0) | 78.7 (1.6) | 65.6 (3.2) |  | 1.3 (1.8%) | 13.1 (16.6%) |
| UP | 39.9 (0.3) | 39.5 (0.9) | 31.3 (2.6) |  | 0.4 (1.1%) | 8.2 (20.8%) |
| UU | 39.9 (0.3) | 39.3 (1.1) | 31.7 (2.1) |  | 0.6 (1.6%) | 7.6 (19.3%) |

**S3 Table.** Results of the cue $\times$ pic RM-ANOVA on ECG amplitudes at R peaks.

| Effect | *Cue* phase | | |  | *Pic* phase | | |
| --- | --- | --- | --- | --- | --- | --- | --- |
|  | *F* | *p* | n_p_^2^ |  | *F* | *p* | n_p_^2^ |
| Cue | 2.247 | .146 | .082 |  | 0.430 | .517 | .017 |
| Pic | 0.762 | .391 | .030 |  | 2.238 | .135 | .087 |
| Cue $\times$ pic | 0.179 | .676 | .007 |  | 0.004 | .952 | .000 |

**S4 Table.** Correlation between BDI scores and ECG amplitudes at R peaks for the UU, PU, UP, and UU conditions at the cue and pic phases. BDI: Beck Depression Inventory; PP: predictably pleasant; PU: predictably unpleasant; UP: unpredictably pleasant; and UU: unpredictably unpleasant.

| Condition | *Cue* phase | | | |  | *Pic* phase | | | |
| --- | --- | --- | --- | --- | --- | --- | --- | --- | --- |
|  | *r* | *p* | *r*^2^ | Power |  | *r* | *p* | *r*^2^ | Power |
| PP | .194 | .364 | .038 | .159 |  | .192 | .369 | .037 | .157 |
| PU | .193 | .365 | .037 | .159 |  | .189 | .376 | .036 | .154 |
| UP | .197 | .357 | .039 | .163 |  | .192 | .368 | .037 | .157 |
| UU | .195 | .362 | .038 | .160 |  | .190 | .375 | .036 | .154 |

**S5 Table.** Correlation between BDI scores and ECG amplitudes averaged over periods for HEP clusters in the UU condition at the cue and pic phases. BDI: Beck Depression Inventory; UU: unpredictably unpleasant.

| Duration | *Cue* phase | | | |  | *Pic* phase | | | |
| --- | --- | --- | --- | --- | --- | --- | --- | --- | --- |
|  | *r* | *p* | *r*^2^ | Power |  | *r* | *p* | *r*^2^ | Power |
| 0.10–0.36 s | .389 | .056 | .156 | .532 |  | .402 | .051 | .162 | .547 |
| 0.10–0.29 s | .379 | .068 | .144 | .495 |  | .398 | .054 | .158 | .537 |
| 0.29–0.33 s | .375 | .071 | .141 | .485 |  | .363 | .082 | .132 | .457 |

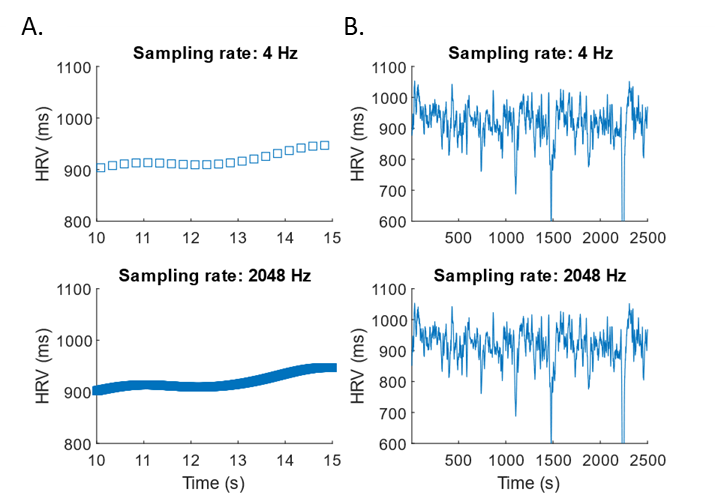

**S1 Figure.** **Comparison of the heart rate variability (HRV) time series before and after upsampling**. HRV time series were upsampled from 4 Hz to 2,048 Hz over (a) a short segment (5 s) and (b) a long segment (2,500 s). The data are from one of the participants. The top and bottom panels show the time series before and after the upsampling process, respectively. In (a), each square represents a single sample of HRV. The squares in the bottom overlap with each other and thus resemble a thick line. These figures show that the HRV after upsampling was similar to that before upsampling.

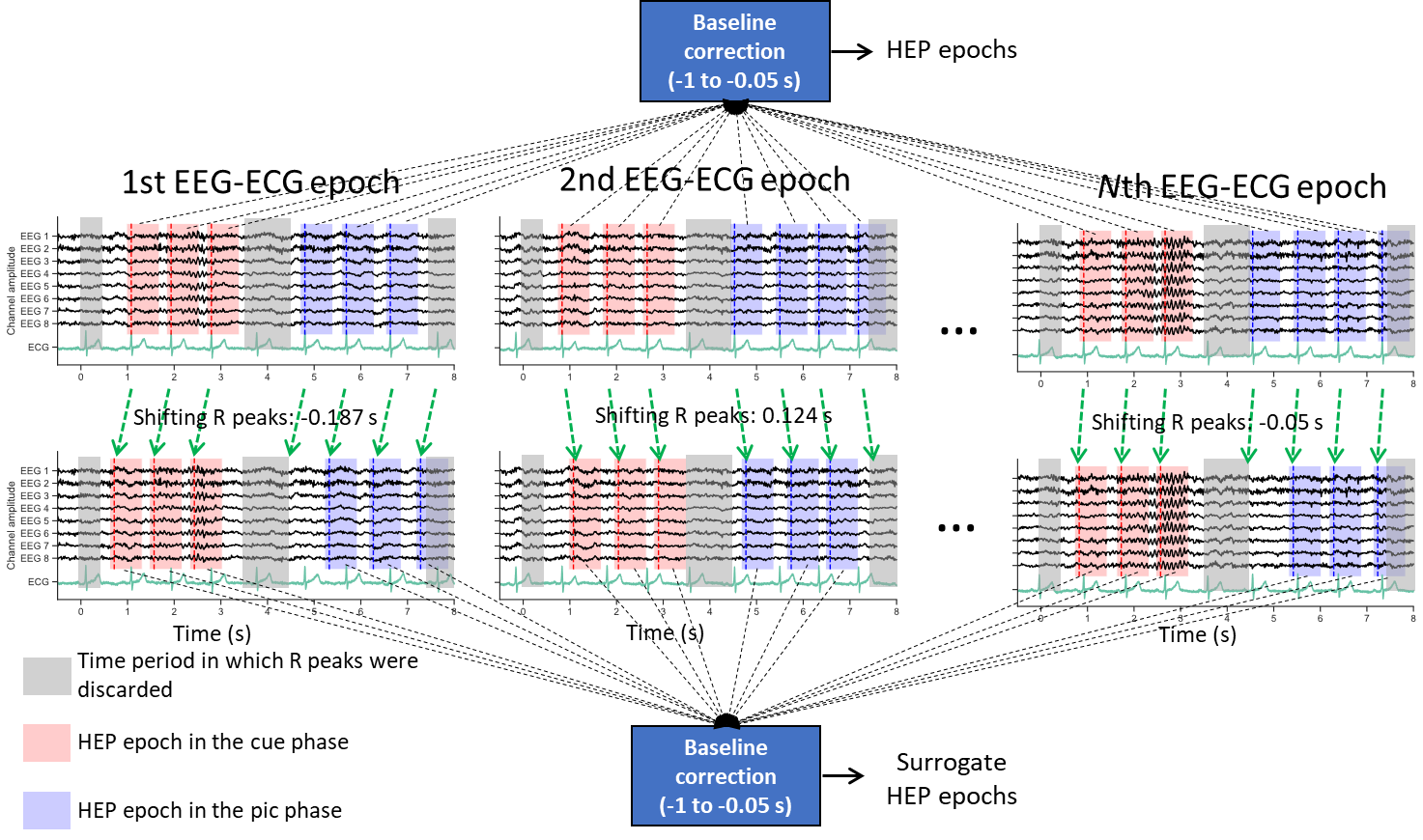
 **S2 Figure.** **Diagram showing how real and surrogate heartbeat-evoked potential (HEP) epochs were calculated.** HEP epochs were obtained by segmenting EEG epochs during the period of -0.1 to 0.6 s with the true R peak time as the onset time, followed by baseline correction. Surrogate R peaks were jittered from true R peaks in the range of -0.5 to 0.5 s (green dashed arrows). Surrogate HEP epochs were then obtained by segmenting EEG epochs during the interval of -0.1 to 0.6 s, with surrogate R peak times as the onset time, followed by baseline correction. Note that the R peaks and surrogate R peaks occurring outside the ranges of 0.5–3.4 s and 4.5–7.4 s were discarded.

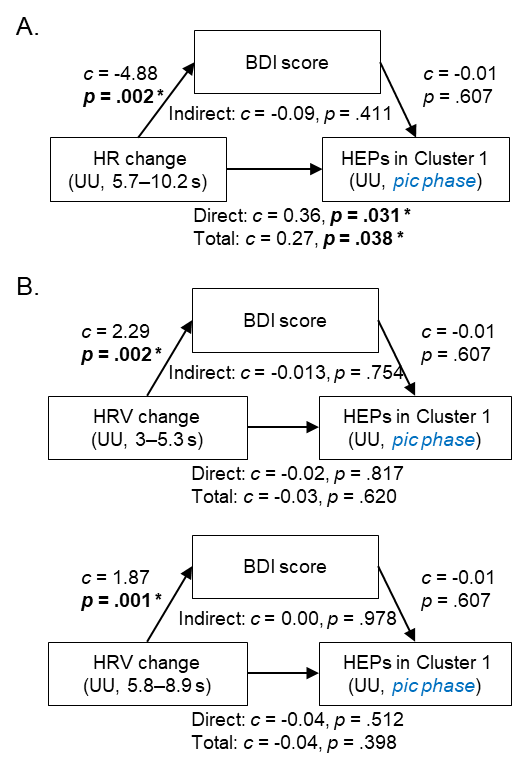

**S3 Figure.** **Results of mediation analysis.** The analysis examining whether depression risk mediates the relationship between cardiac reactivity, reflected by (a) heart rate (HR) and (b) heart rate variability (HRV), and neural responses to emotional pictures, represented by heartbeat-evoked potentials (HEPs) in the picture (pic) phase. No significant mediation effect was found in these three models. BDI: Beck Depression Inventory; UU: unpredictably unpleasant.

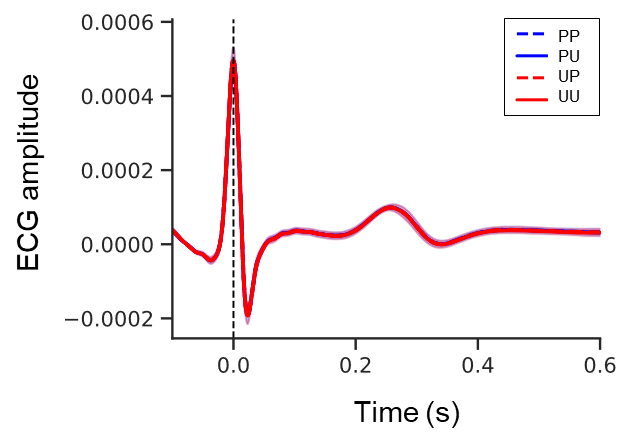

**S4 Figure.** **Time series of averaged ECG epochs for the PP, PU, UP and UU conditions.** The vertical black dashed line indicates the time of the R peaks. PP: predictably pleasant; PU: predictably unpleasant; UP: unpredictably pleasant; and UU: unpredictably unpleasant.

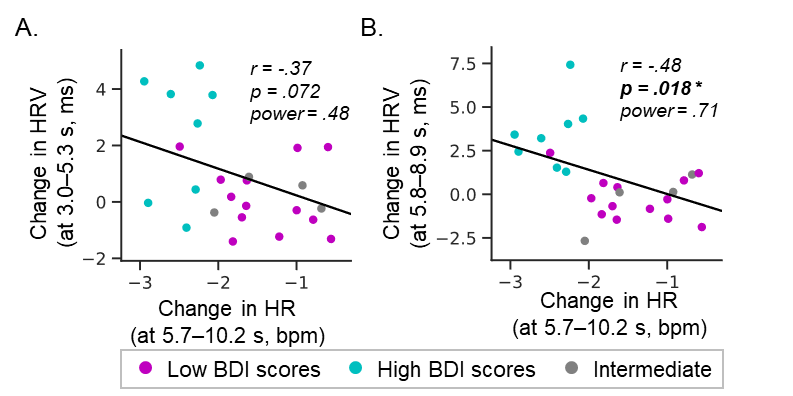

**S5 Figure.** **Scatter plots of changes in heart rate (HR) and heart rate variability (HRV).** The changes in HR were averaged over the interval of 5.7–10.2 s and those in heart rate variability (HRV) were averaged over the intervals of (a) 3–5.3 s and (b) 5.8–8.9 s in the unpredictably unpleasant (UU) condition. Black solid lines represent regression fits of data from all participants.
